# The joint action of antibiotics, bacteriophage, and the innate immune response in the treatment of bacterial infections

**DOI:** 10.1101/2025.04.10.648192

**Authors:** Brandon A. Berryhill, Teresa Gil-Gil, Bruce R. Levin

## Abstract

Studies of antimicrobial therapeutics have traditionally neglected the contribution of the host in determining the course of treatment and its outcome. One critical host element, which shapes the dynamics of treatment is the innate immune system. Studies of chemotherapeutics and complementary therapies such as bacteriophage (phage), are commonly performed with mice that purposely have an ablated innate immune system. Here, we generate a mathematical and computer-simulation model of the joint action of antibiotics, phage, and phagocytes. Our analysis of this model highlights the need for future studies to consider the role of the host’s innate immune system in determining treatment outcomes. Critically, our model predicts that the conditions under which resistance to the treatment agent(s) will emerge are much narrower than commonly anticipate. We also generate a second model to predict the dynamics of treatment when multiple phages are used. This model provides support for the application of cocktails to treat infections rather than individual phages. Overall, this study provides hypotheses that can readily be tested experimentally with both *in vitro* and *in vivo* experiments.

## Introduction

As a consequence of the increasing frequency of infections with antibiotic resistant bacteria there has been an increase in research on and the application of bacteriophage (phage) for the treatment of bacterial infections (1, 2). Phage therapy was employed before the advent of antibiotics but was ultimately replaced by these drugs; however, the recent resurrection of phage therapy sees these viruses used concomitantly with antibiotics almost exclusively (3–5). There is currently a lack of understanding about the interactions between these bacterial viruses and these drugs, especially in regards to the conditions where they act either synergistically or antagonistically (6). The purpose of this this report is to use mathematical and computer simulation models to explore the population dynamic and evolutionary processes required for effective therapy with antibiotics and phage.

In exploring joint phage and antibiotic therapy, it is critical to consider the contribution of the host’s innate immune system in the control of bacterial infections. The innate immune defenses play a prominent role to the course of antibiotic and phage therapy and need to be considered in studies evaluating the effect of these agents both independently and when used together (7–9). In this report, we restrict our consideration of the innate immune system to phagocytes and phagocytosis. We give particular focus to the effect that dosing (e.g., antibiotic first, phage first, or coadministration) has to the outcomes of infection treatment and the role that the emergence of antibiotic and phage resistances have on treatment dynamics.

Phages, when given to treat an infection, are often administered as cocktails of multiple phages, with one of the primarily goals being preventing the ascent of phage resistance, which would decrease treatment efficacy (10). The majority of our modeling results presented here assume only one phage and antibiotic are used; however, we do employ a second model to determine the contribution multiple phages would have to the dynamics of treated bacterial given the emergence of phage resistant mutants.

## Mathematical models

### A model of phage, antibiotics, and the innate immune system

The model developed here is an extension of that in (11) which has been expanded to include phage. This model assumes continuous culture (chemostat) conditions (12). Shown in Figure 1 is a diagram of the model employed in this report. Tables 1 and 2 detail the variables of this model and the default parameters used in our simulations, respectively.

**Figure 1.**
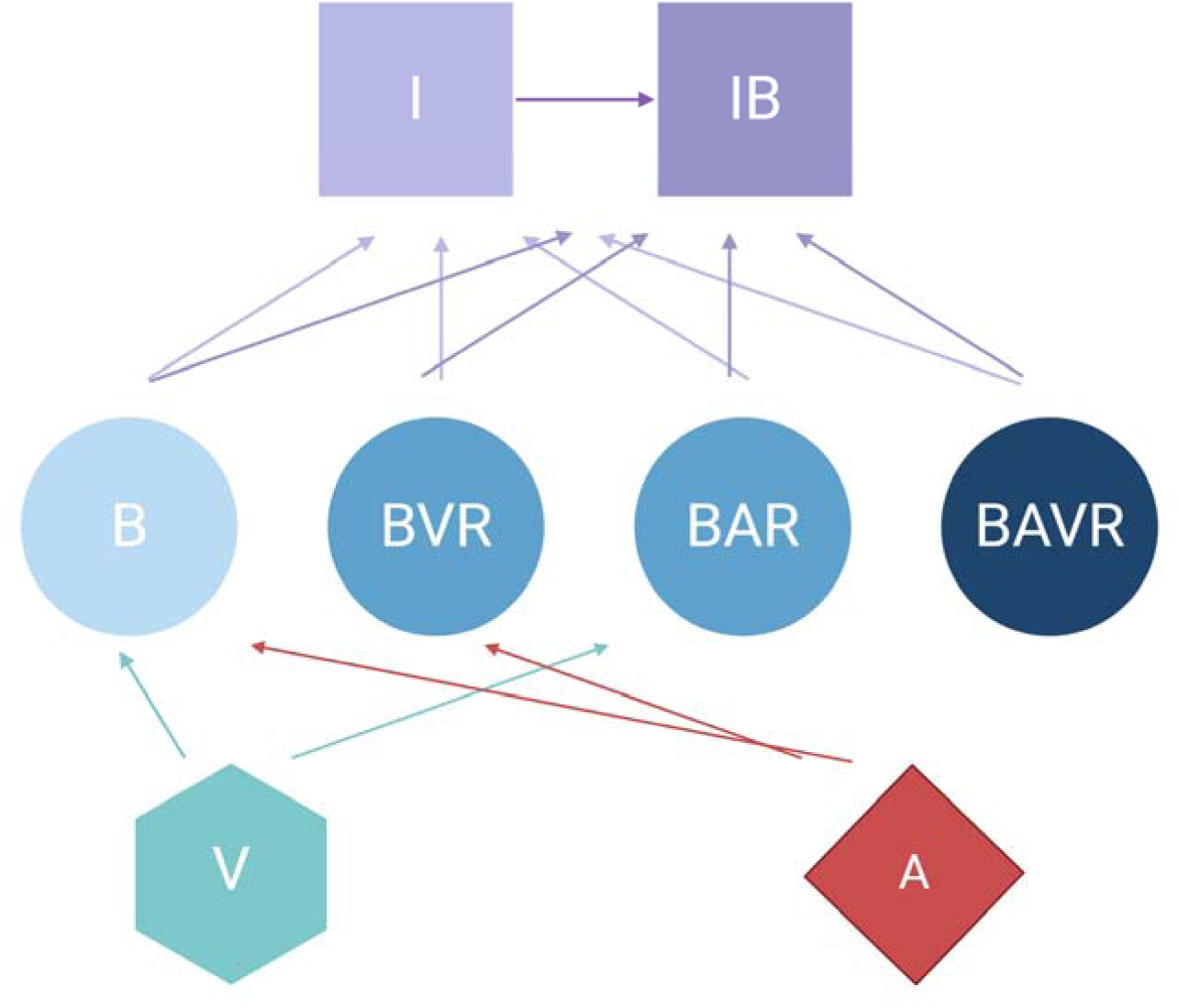
Diagram of the model of the joint action of antibiotics, phage, and the innate immune system in the dynamics of treatment of a bacterial infection. For the definitions of the variable in the above diagram, see Table 1. All parameters, their definitions, and values used in the simulations of this model are presented in Table 2 unless otherwise stated.

**Table 1.** Variables used in the model of the joint action of antibiotics, phage, and the innate immune system.

|  |  |
| --- | --- |
| R | The limiting resource |
| B | Antibiotic-sensitive, phage-sensitive bacteria |
| BVR | Antibiotic-sensitive, phage-resistant bacteria |
| BAR | Antibiotic-resistant, phage-sensitive bacteria |
| BAVR | Antibiotic-resistant, phage-resistant bacteria |
| V | Lytic phage |
| A | Antibiotic |
| I | Free phagocytes |
| IB | Phagocytes that have engulfed at least one bacterium |

**Table 2.** Parameters and the values used in the simulation of the joint action of antibiotics, phage, and the innate immune system.

| Parameter | Definition | Value | Units | Source |
| --- | --- | --- | --- | --- |
| $vb, vbar, vbvr, vbavr$ | Maximum growth rates | 1.0, 0.9, 0.9, 0.8 | per cell per hour | This report |
| C | Maximum resource concentration | 1000 | $\mu\text{g/mL}$ | This report |
| e | Resource conversion efficiency | $5 \cdot 10^{-7}$ | $\mu\text{g/cell}$ | (13) |
| $\mu$ | Mutation rate | $10^{-7}$ | per hour | (14) |
| IMAX | Maximum phagocyte density | $10^5$ | per mL | This report |
| $\gamma$ | Phagocyte gobbling constant | $9 \cdot 10^{-6}$ | per cell per hour | This report |
| w | Flow rate | 0.1 | mL per hour | (12) |
| $\delta$ | Phage adsorption rate | $10^{-7}$ | per hour per mL | (15) |
| $\beta$ | Phage burst size | 50 | particles per cell | (15) |
| da | Decay rate of the antibiotic | 0 | $\mu\text{g/mL}$ per hour | (16) |
| $vmin$ | Maximum kill rate of antibiotic | -4.0 or -0.001 | per cell per hour | (17) |
| $\kappa$ | Hill parameter | 1.0 | | (18) |
| MIC | Minimum inhibitory concentration | 1.0 | $\mu\text{g/mL}$ | This report |
| k | Monod constant | 1.0 | $\mu\text{g}$ | (19) |
| Dose | Dosing interval | 8.0 | Hours | This report |

#### Pharmacodynamics of antibiotic treatment

We assume that resource (*R*, µg/mL) enters the environment at a constant rate and that the pharmacodynamics of the antibiotics and bacteria are modeled by a Hill function where: ∏ *(A, R)* is the net growth/death rate of the bacteria (Equation 1) (18). Equation 1 is written generally with *i* in place of the bacterial states (e.g., B or BVR). For this model, we assume the resource is the unique agent limiting the growth and final density in the absence of antibiotics, phage, or the immune system—analogous to the carbon source in a minimal media (13). To simulate the effect that the decreasing limiting resource concentration has on the physiological state of the bacteria, we include a term ψ(R) defined in Equation 2 (19).

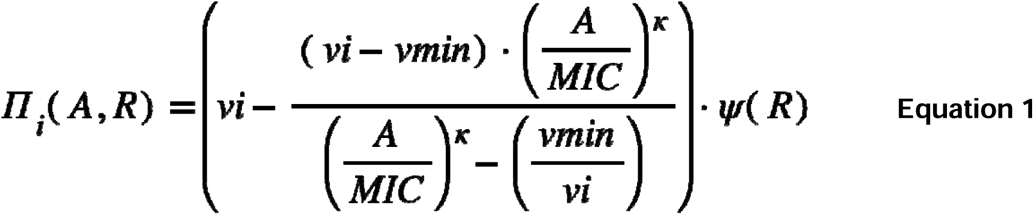

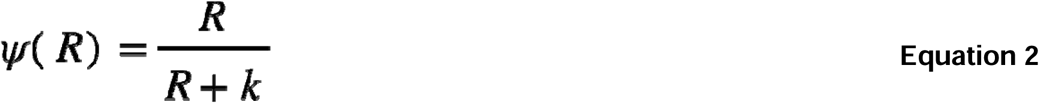

#### Mathematical model of phage, antibiotics, and the innate immune system

To simulate the treatment of populations of antibiotic-sensitive and phage-sensitive bacteria, antibiotic-resistant and phage-sensitive bacteria, antibiotic-sensitive and phage-resistant bacteria, and antibiotic-resistant and phage-resistant bacteria with antibiotics and phage, we construct a series of coupled, ordered differential equations (Equations 3 through 11). With these definitions, assumptions, and the parameters defined and presented in Tables 1 and 2, the rates of change in the densities of the different populations are given by:

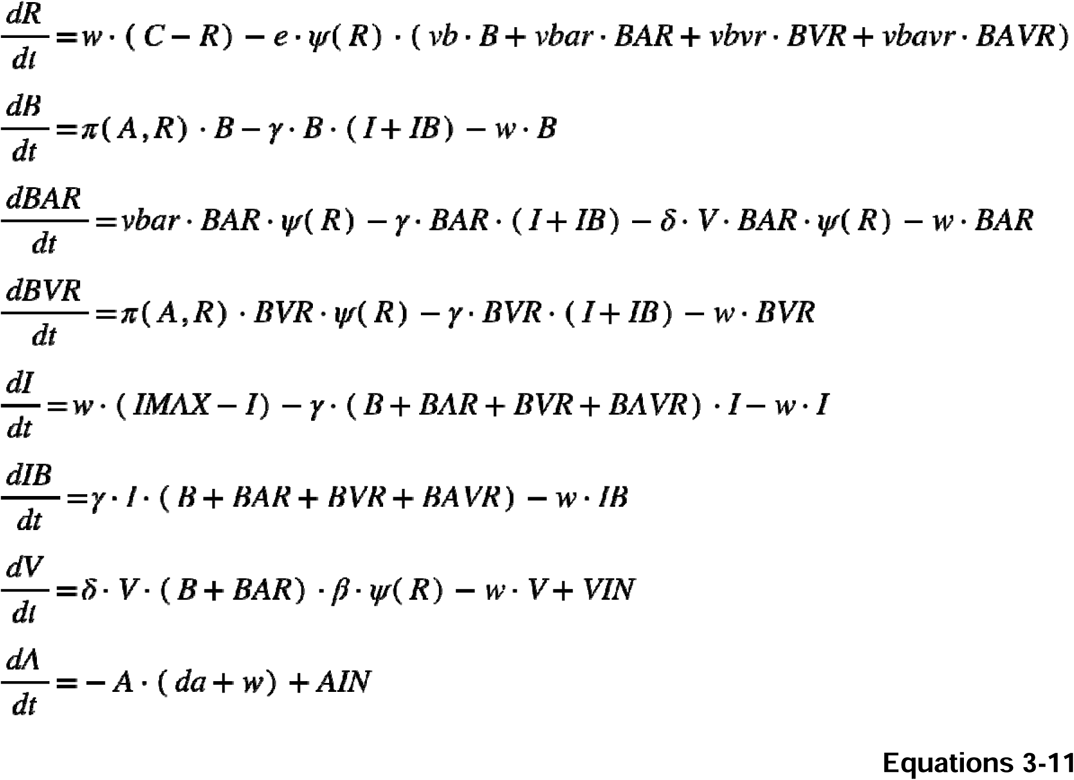

### A model of phage cocktails

Shown in Figure 2 is a diagram of the model employed for modeling phage cocktails in this report. Tables 3 and 4 detail the variables of this model and the default parameters used in our simulations, respectively.

**Figure 2.**
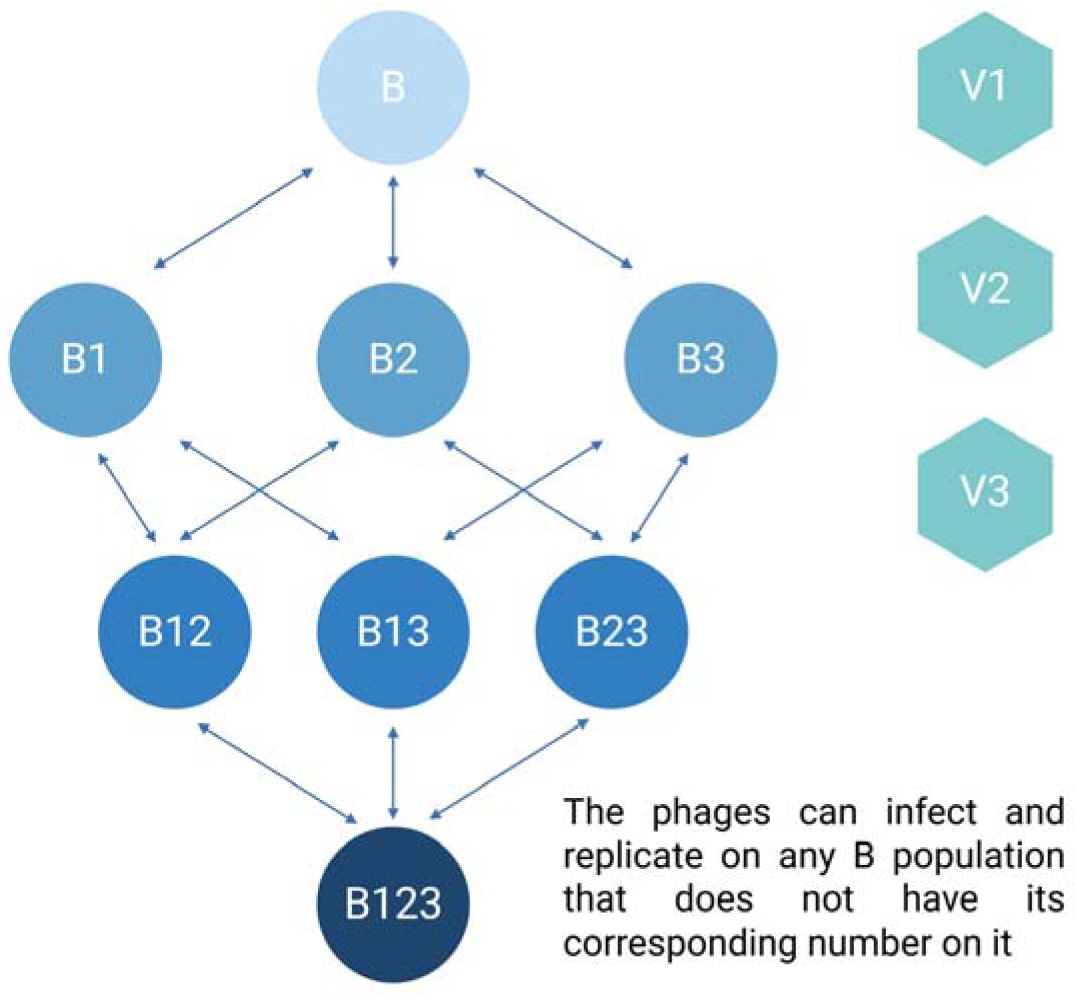
Diagram of the phage cocktail model. For the definitions of the variables in the above diagram, see Table 3. All parameters, their definitions, and values used in the simulations of this model are presented in Table 4 unless otherwise stated.

**Table 3.** Variables used in the model of bacteriophage cocktails.

|  |  |
| --- | --- |
| R | The limiting resource |
| V1 | Phage 1 |
| V2 | Phage 2 |
| V3 | Phage 3 |
| B | Bacteria susceptible to all phages |
| BR1 | Bacteria resistant to phage 1 |
| BR2 | Bacteria resistant to phage 2 |
| BR3 | Bacteria resistant to phage 3 |
| BR12 | Bacteria resistant to phage 1 and phage 2 |
| BR13 | Bacteria resistant to phage 1 and phage 3 |
| BR23 | Bacteria resistant to phage 2 and phage 3 |
| BR123 | Bacteria resistant to phage 1, phage 2, and phage 3 |

**Table 4.**
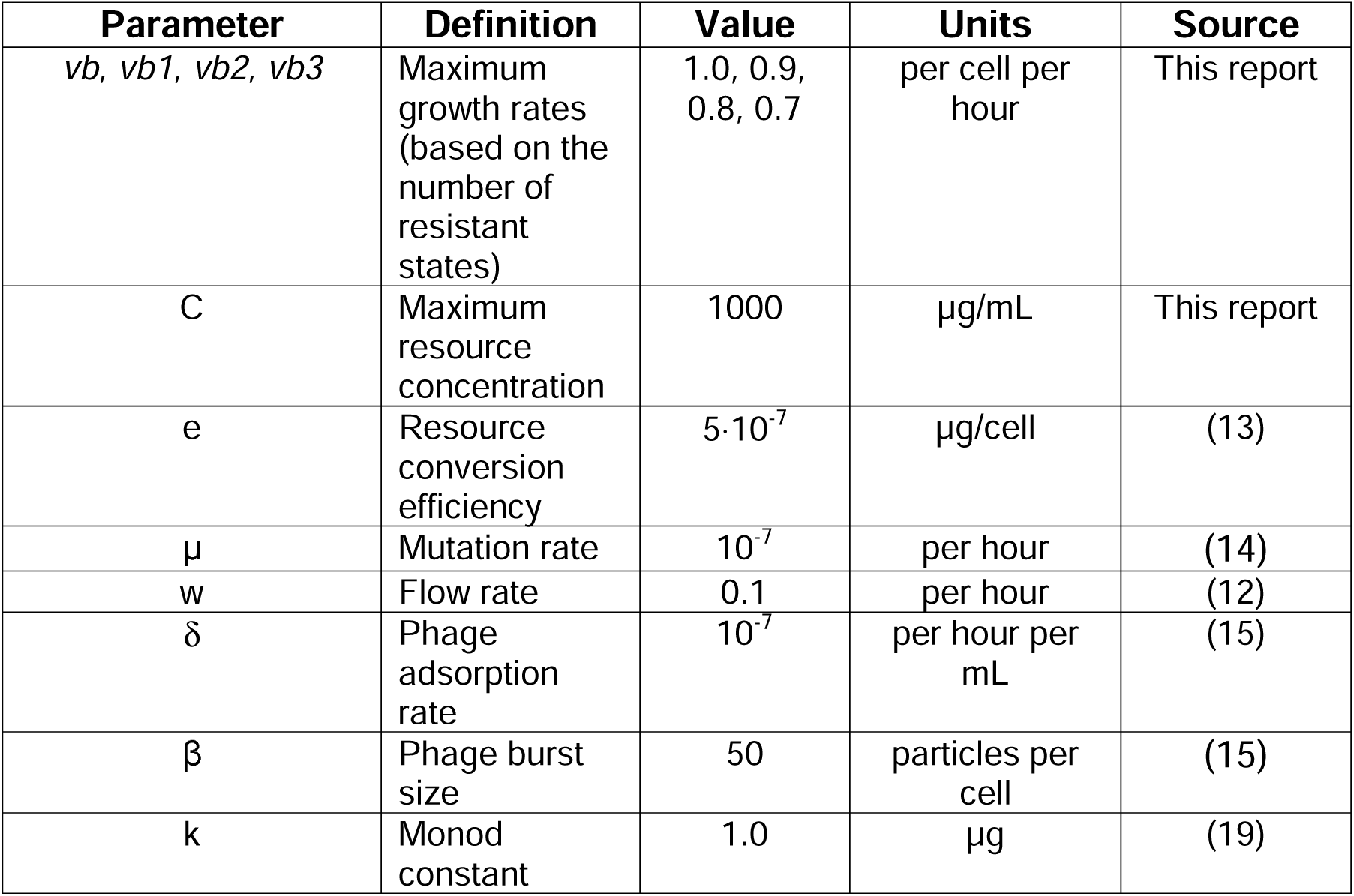
Parameters and the values used in the model of bacteriophage cocktails.

| Parameter | Definition | Value | Units | Source |
| --- | --- | --- | --- | --- |
| $vb, vb1, vb2, vb3$ | Maximum growth rates (based on the number of resistant states) | 1.0, 0.9, 0.8, 0.7 | per cell per hour | This report |
| C | Maximum resource concentration | 1000 | $\mu\text{g/mL}$ | This report |
| e | Resource conversion efficiency | $5 \cdot 10^{-7}$ | $\mu\text{g/cell}$ | (13) |
| $\mu$ | Mutation rate | $10^{-7}$ | per hour | (14) |
| w | Flow rate | 0.1 | per hour | (12) |
| $\delta$ | Phage adsorption rate | $10^{-7}$ | per hour per mL | (15) |
| $\beta$ | Phage burst size | 50 | particles per cell | (15) |
| k | Monod constant | 1.0 | $\mu\text{g}$ | (19) |

#### Mathematical model of phage cocktails

To simulate the treatment of populations of bacteria which are either phage-sensitive, resistant to one phage, resistant to two phages, or resistant to three phages and three different phages we construct a series of coupled, ordered differential equations (Equations 12 through 23). In this model, the transitions between the various phage-resistant states occurs stochastically (20). We simulate these transitions with a Monte Carlo process (21). A random number x (0 ≤ x ≤1) from a rectangular distribution is generated (22). If x is less than the product of the number of cells in the generating state (B, the density time the volume of the vessel, Vol), the transition rate (µ) and the step size (dt) of the Euler method employed for solving the differential equations (23), for example if x < B*µ*dt*Vol, then ADDBR1B cells are added to the BR1 population and removed from the B population where ADDBR1B=1/(dt*Vol). With these definitions, assumptions, and the parameters defined and presented in Tables 3 and 4, the rates of change in the densities of the different populations are given by:

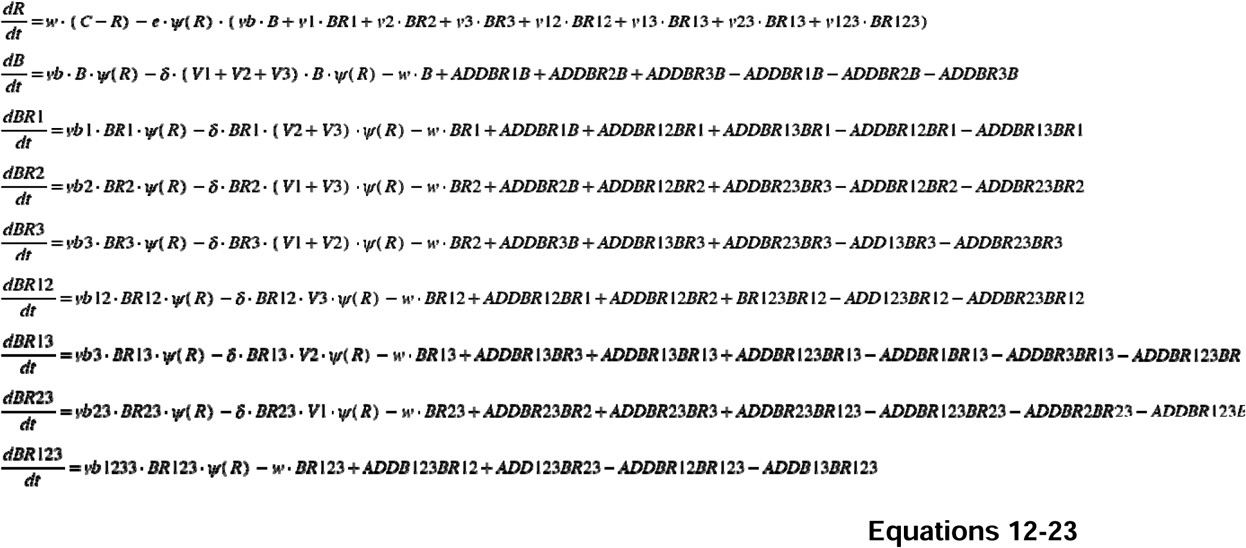

## Results

### Control by the immune system in the absence of treatment

We begin our analysis of the predictions generated by the first model by considering the effect the primary, unmeasured parameter γ (the phagocyte gobbling rate) has on the dynamics of the infection. In Figure 3, we consider three differing values of γ, demonstrating that the model is highly sensitive to this parameter. Going forward, all simulations are performed with the value of γ in Figure 3A, where the immune system is capable of suppressing the growth of the bacteria but not capable of clearing the infection on its own.

**Figure 3.**
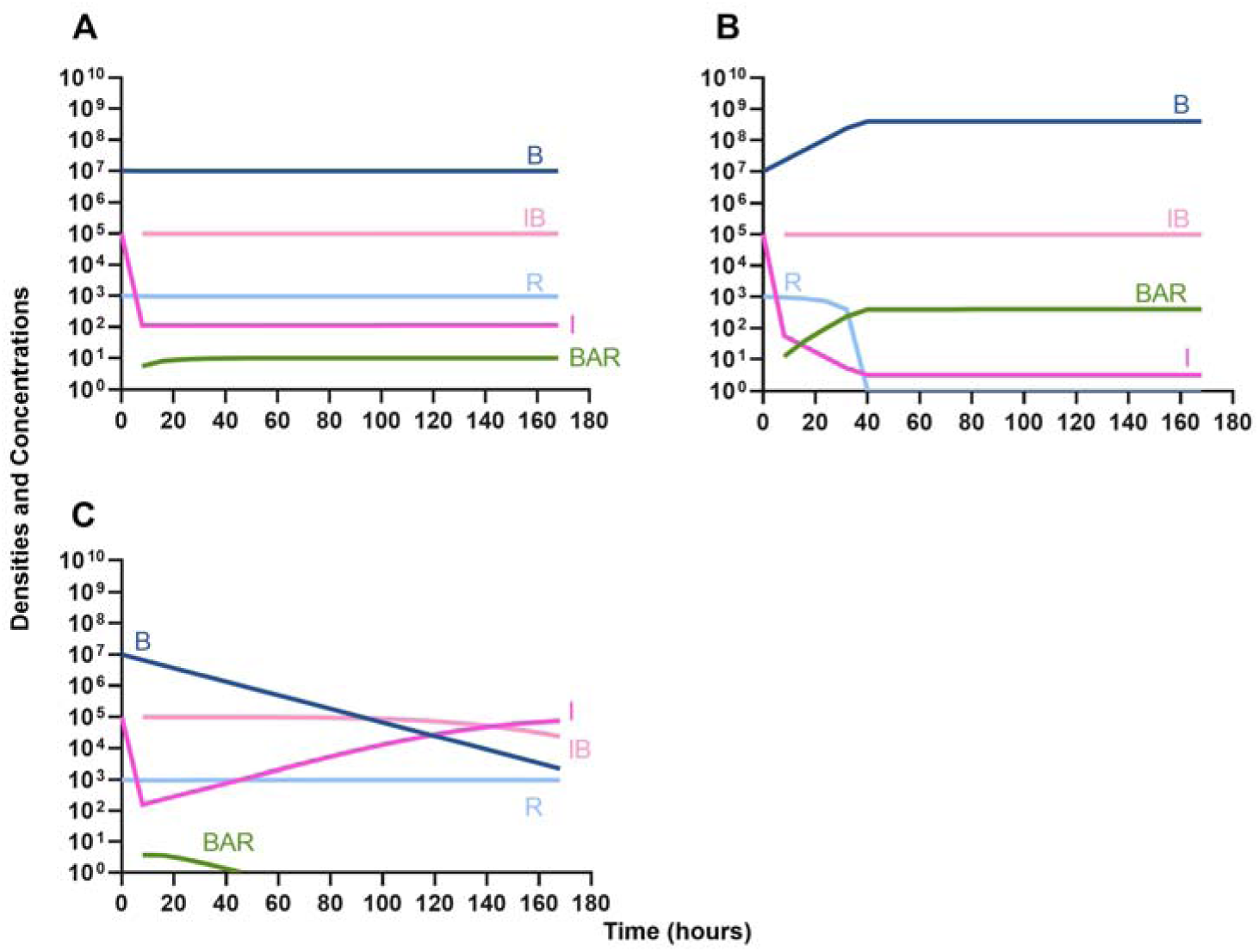
Effect of immune system on controlling infections without treatment. **A.** The level of immune response needed to prevent net growth or death of the bacteria populations. γ = 9E-6 **B.** A marginally weaker immune response. γ = 8E-6 **C.** A marginally stronger immune response. γ = 9.5E-6.

### Treatment of infections in the absence of the immune system

We then analyze the effects that treatment in the absence of the innate immune system has on the dynamics of infection.

#### Single agent treatment (controls)

In Figure 4A we consider treatment of an initially sensitive bacteria population with a highly bactericidal antibiotic; in Figure 4B the only treatment is a bacteriostatic drug; and, in Figure 4C the bacteria are treated with a lytic bacteriophage. Notably, all agents are capable of controlling the initial infection, however resistance to that agent does rapidly ascend.

**Figure 4.**
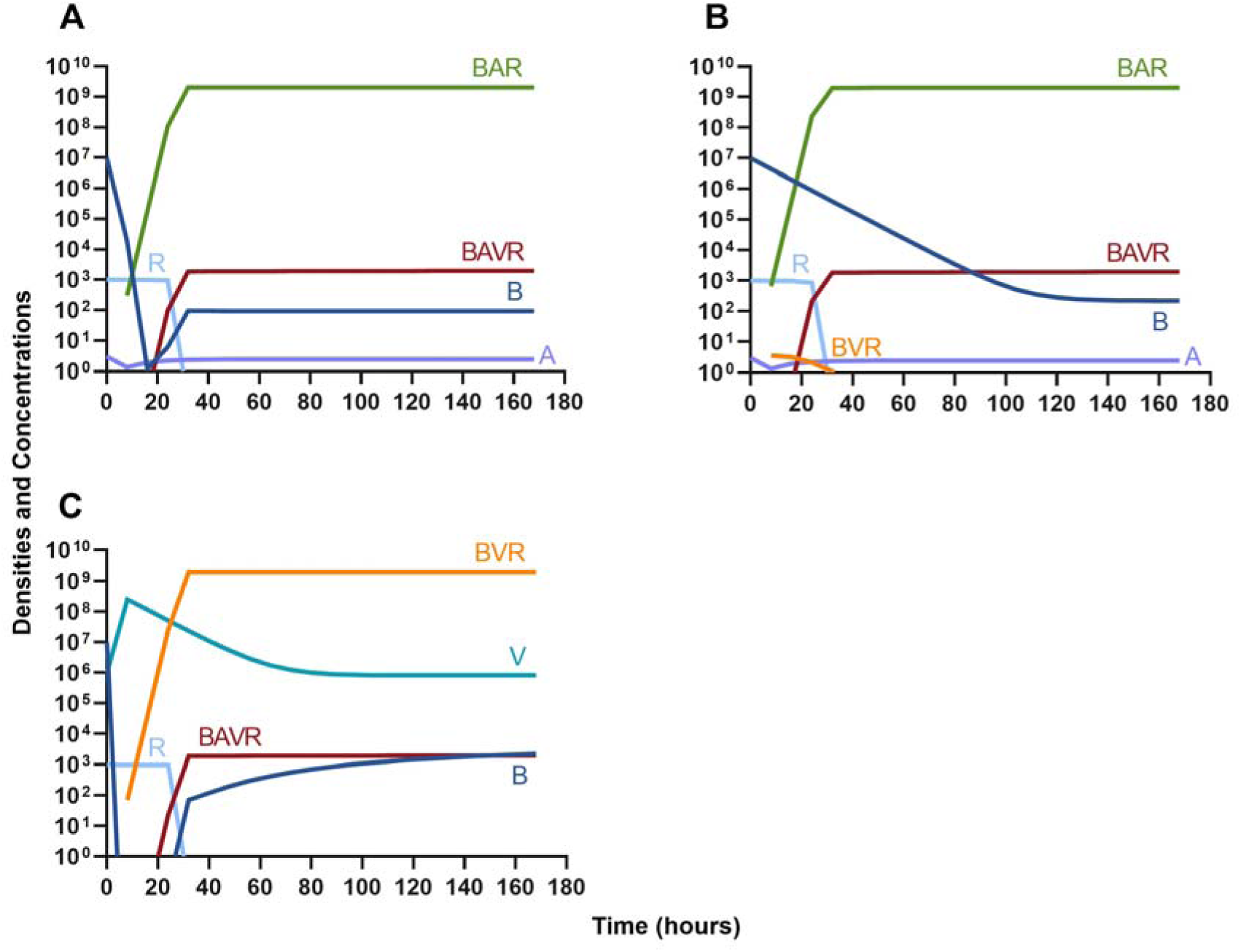
Single agent treatment without the immune system. **A.** A bactericidal antibiotic **B.** A bacteriostatic antibiotic **C.** A lytic bacteriophage.

#### Bacteriostatic antibiotics

In evaluating the joint action of phage and antibiotics, we first examine a highly lytic phage in combination with a bacteriostatic antibiotic. There are three distinct dosing regimens: phage first (Figure 5A), bacteriostatic drug first (Figure 5B), and coadministration of both the phage and antibiotic (Figure 5C). Our simulations predict that the phage first regimen clears the initial bacterial population the quickest, followed by antibiotic first, and then coadministration being the slowest to clear the initial population. However, in all cases resistance to both treating agents ascends in roughly the same amount of time, approximately 40 hours.

**Figure 5.**
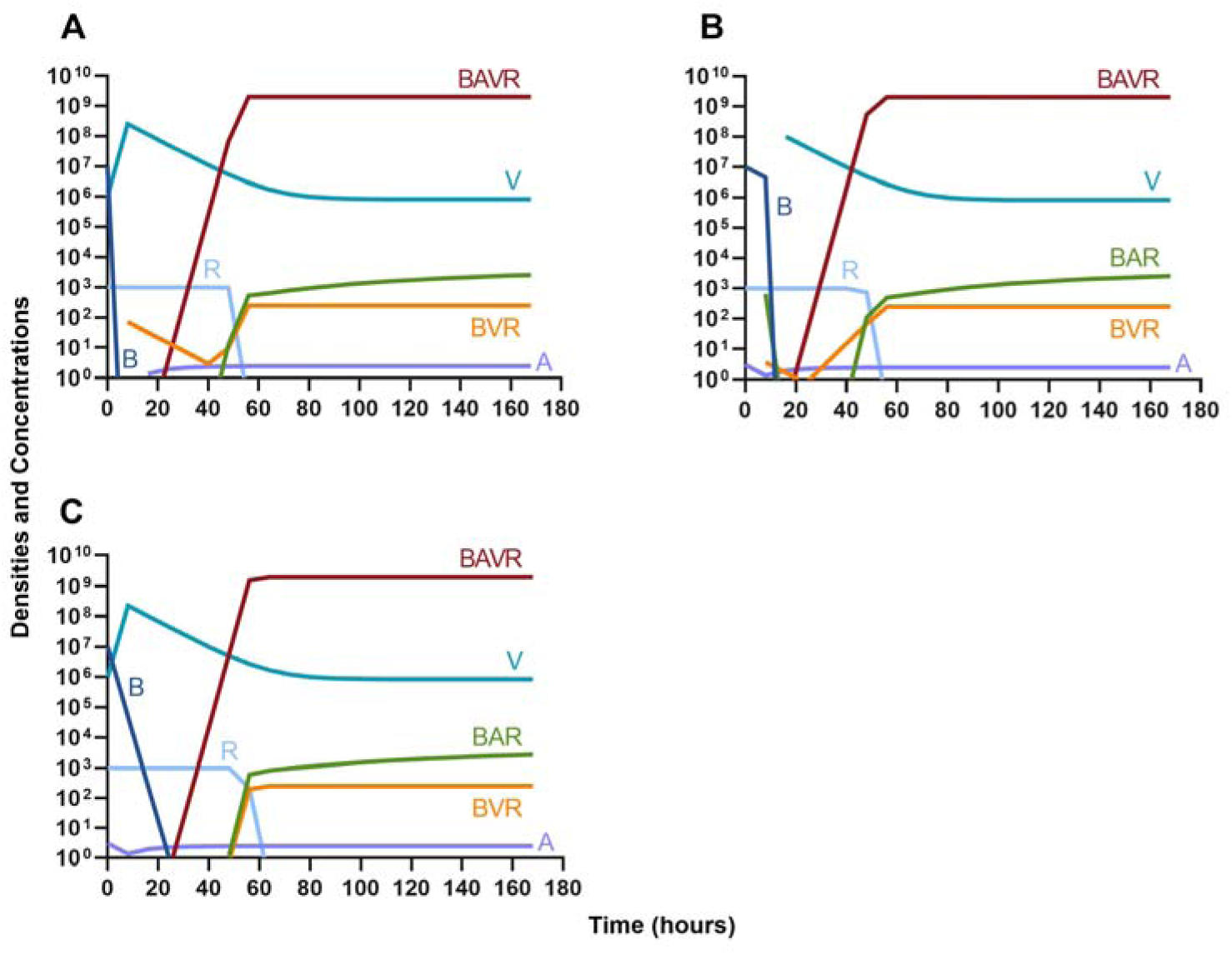
Treatment with a bacteriostatic drug and a bacteriophage with differing dosing regimens. **A.** A phage and then a bacteriostatic antibiotic **B.** A bacteriostatic antibiotic and then a phage **C.** Coadministration of a bacteriostatic antibiotic and a phage.

#### Bactericidal antibiotics

We continue our investigation of the dynamics of treatment without the immune system by studying the joint action of a bactericidal drug and a phage. Again, there are three distinct dosing regimens: phage first (Figure 6A), bactericidal drug first (Figure 6B), and coadministration of both the phage and antibiotic (Figure 6C). Our simulations provide the same predictions as those for the bacteriostatic drug. Indicating, that both bacteriostatic and bactericidal antibiotics can be equally as effective, however resistant will ultimately ascend.

**Figure 6.**
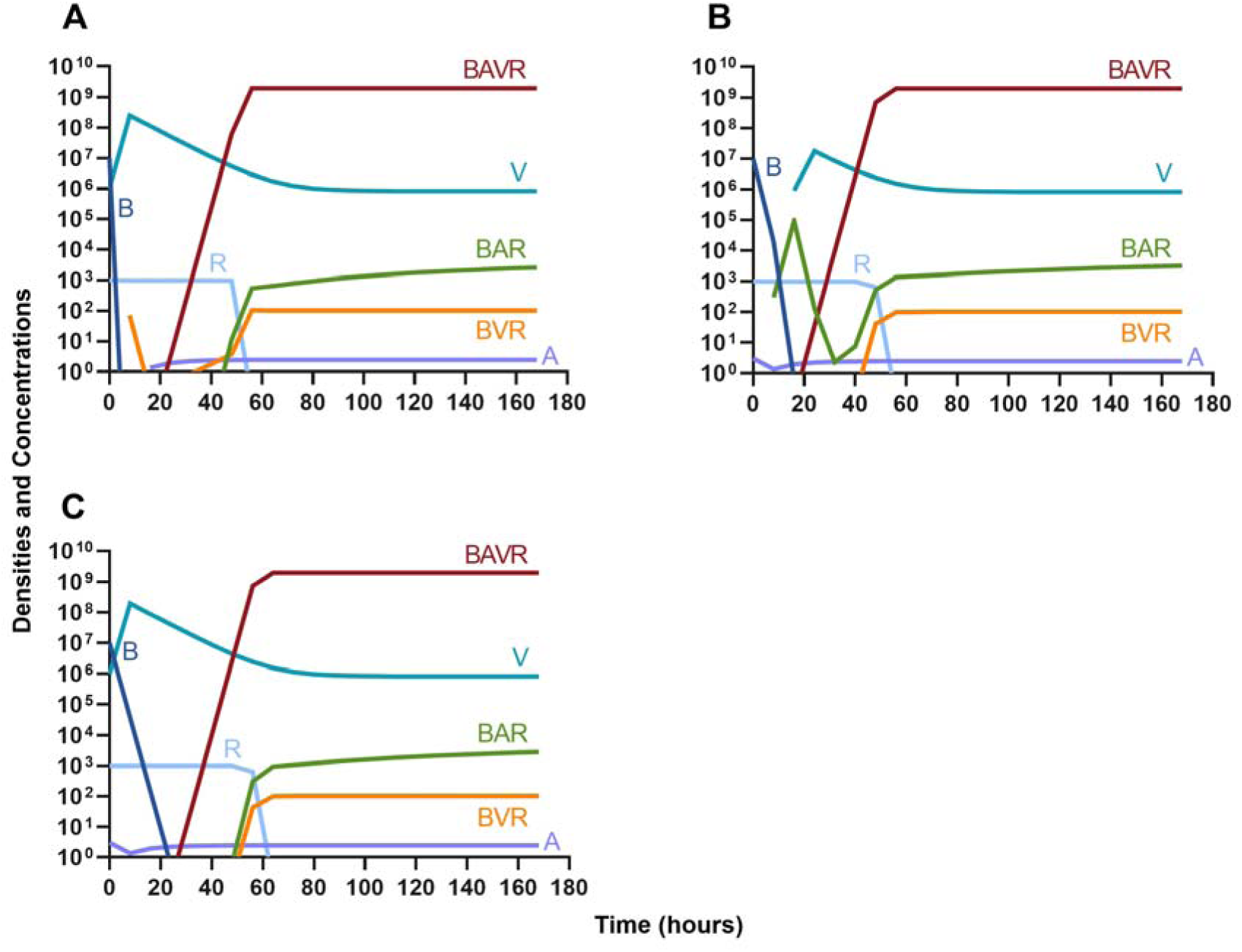
Treatment with a bactericidal drug and a bacteriophage with differing dosing regimens. **A.** A phage and then a bactericidal antibiotic **B.** A bactericidal antibiotic and then a phage **C.** Coadministration of a bactericidal antibiotic and a phage.

#### Phage suppression of antibiotic resistance

One argument for the use of phages in combination with antibiotics is that the virus is able to suppress the antibiotic-resistant population. To address this hypothesis, in Figure 7 we consider a scenario where the majority of the bacteria are susceptible to an antibiotic, but there is a minor population at a ratio of 1:1000 which is resistant to the treating drug (either a bacteriostatic drug as in Figure 7A and 7C or a bactericidal drug Figure 7B and 7D). When the phage is not present (Figure 7A and 7B) the antibiotic-resistant minority population is able to ascend to dominance and treatment fails. While, when the phage is present, the antibiotic-resistant population is rapidly controlled, but a population which is resistant to both the phage and antibiotic ascends to dominance; however, the emergence of this double-resistant population takes twice as long to ascend to dominance.

**Figure 7.**
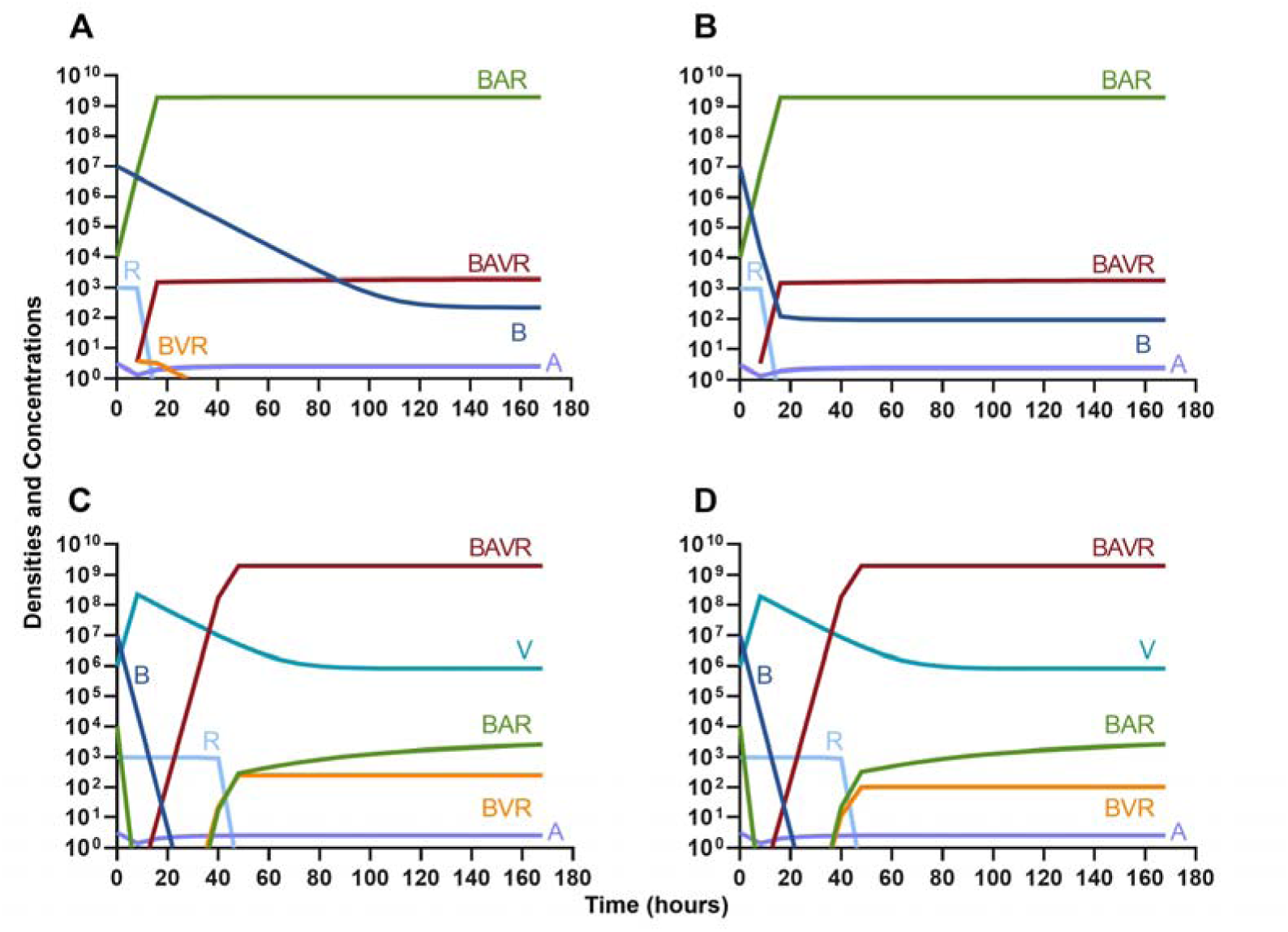
Invasion when rare of antibiotic-resistant bacteria. **A.** A population of 1E7 antibiotic-sensitive bacteria and 1E4 antibiotic-resistant bacteria treated with a bacteriostatic antibiotic **B.** A population of 1E7 antibiotic-sensitive bacteria and 1E4 antibiotic-resistant bacteria treated with a bactericidal antibiotic **C.** A population of 1E7 antibiotic-sensitive bacteria and 1E4 antibiotic-resistant bacteria treated with a bacteriostatic antibiotic and bacteriophage. **D.** A population of 1E7 antibiotic-sensitive bacteria and 1E4 antibiotic-resistant bacteria treated with a bactericidal antibiotic and a bacteriophage.

#### Antibiotic suppression of phage resistance

Finally, we address the same situation as in Figure 7 but instead consider that a phage-resistant population is the minor population present at the initiation of treatment. As expected, in Figure 8A, when treated with just the phage, the phage-resistant population ascends to a majority. As in Figure 7, when treated with either a bacteriostatic (Figure 8B) or bactericidal antibiotic (Figure 8C) in conjunction with the phage, the initial phage-resistant population is controlled, but a population resistant to both treating agents emerges. In this case, the double-resistant mutant takes approximately four times as long to dominate as the single-resistant population.

**Figure 8.**
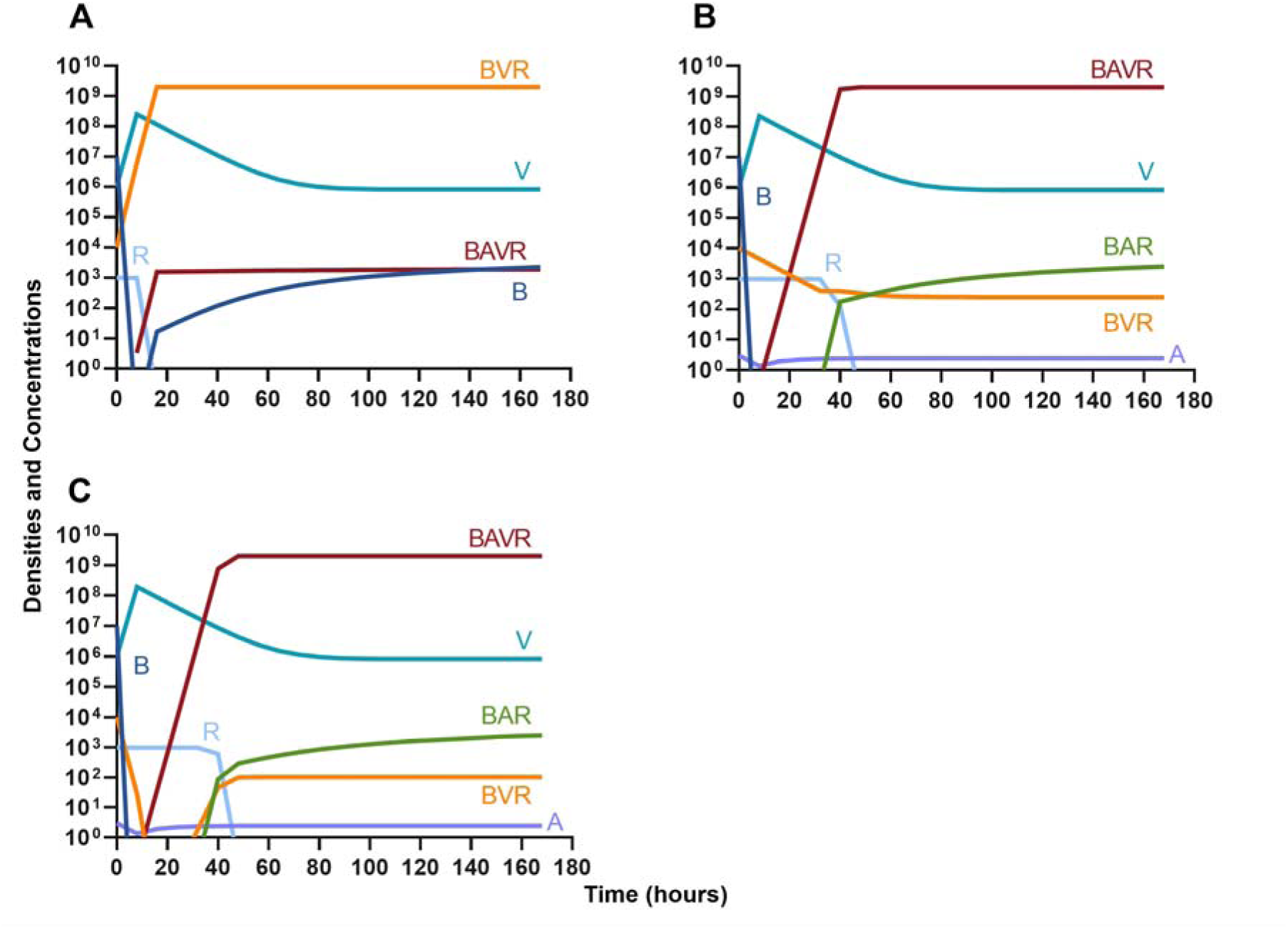
Invasion when rare of phage-resistant bacteria. **A.** A population of 1E7 phage-sensitive bacteria and 1E4 phage-resistant bacteria treated with a bacteriophage **B.** A population of 1E7 phage-sensitive bacteria and 1E4 phage-resistant bacteria treated with a bacteriostatic antibiotic and a phage **C.** A population of 1E7 phage-sensitive bacteria and 1E4 phage-resistant bacteria treated with a bactericidal antibiotic and a phage.

### Single agent treatment and the innate immune system

Given the above results where we do not consider the impact of the innate immune system, we continue our modelling by considering similar situations but with the immune response.

#### Antibiotics and the innate immune system

Given the consideration of the immune system alone in Figure 1, we begin this section, by considering the interaction of antibiotics and the immune system. With both bacteriostatic and bactericidal drugs (Figure 9A and 9B, respectively), the immune system and the antibiotics together rapidly clear the infection, and antibiotic-resistant populations do not appear. Moreover, if the antibiotic-resistant populations are present initially (Figure 9C and 9D), they are rapidly lost as well.

**Figure 9.**
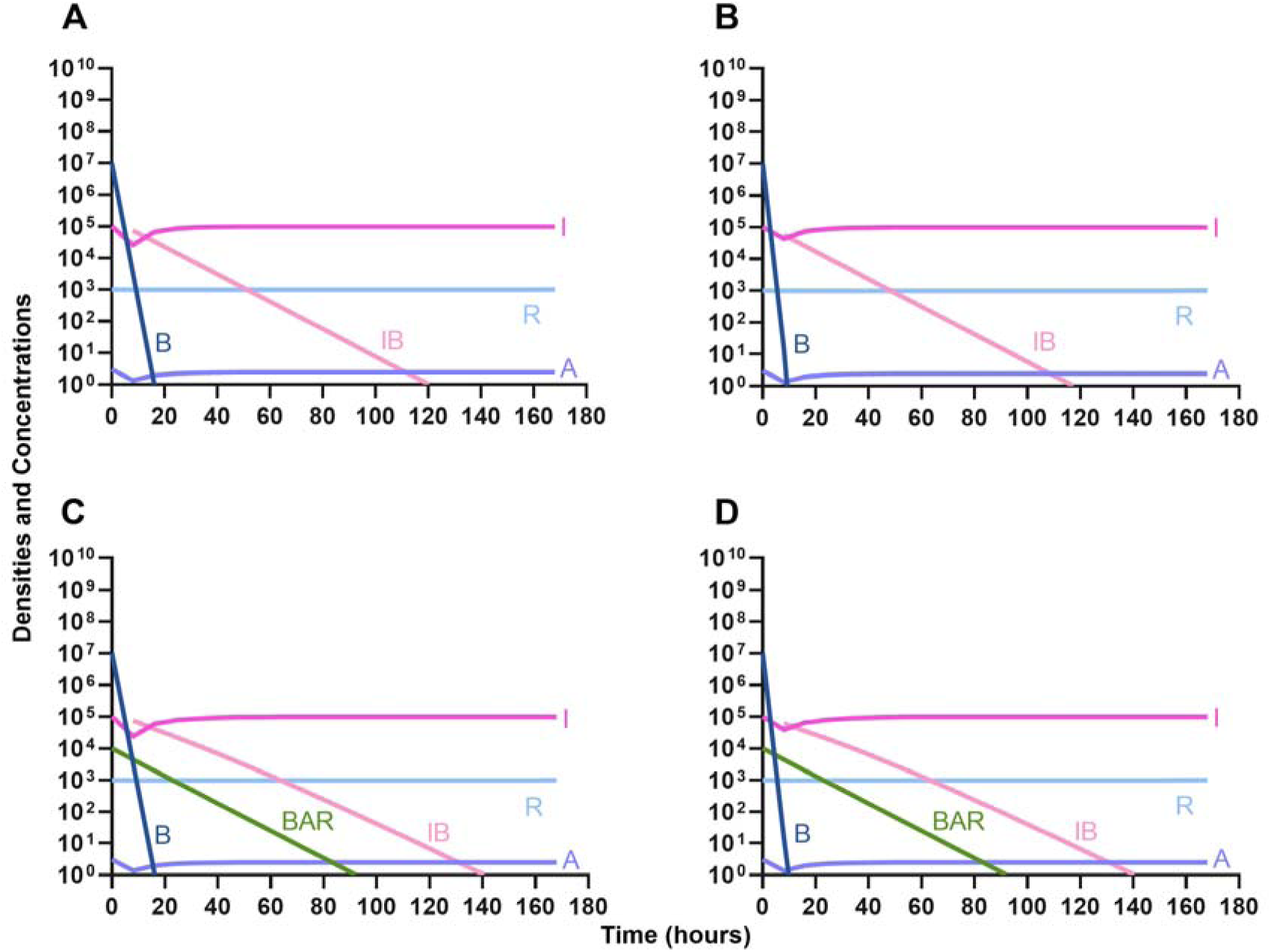
The interaction of antibiotics and the innate immune system. **A.** A population of 1E7 bacteria treated with a bacteriostatic drug in the presence of the immune system **B.** A population of 1E7 bacteria treated with a bactericidal drug in the presence of the immune system **C.** A population of 1E7 bacteria and 1E4 antibiotic-resistant bacteria treated with a bacteriostatic drug in the presence of the immune system **D.** A population of 1E7 bacteria and 1E4 antibiotic-resistant bacteria treated with a bactericidal drug in the presence of the immune system.

#### Phage and the innate immune system

In Figure 10, we consider the same situation as in Figure 9 but instead treat with a lytic phage. As in the previous section, the phage and the immune system can rapidly clear the infection (Figure 10A) and minor phage-resistant populations do not ascend (Figure 10B).

**Figure 10.**
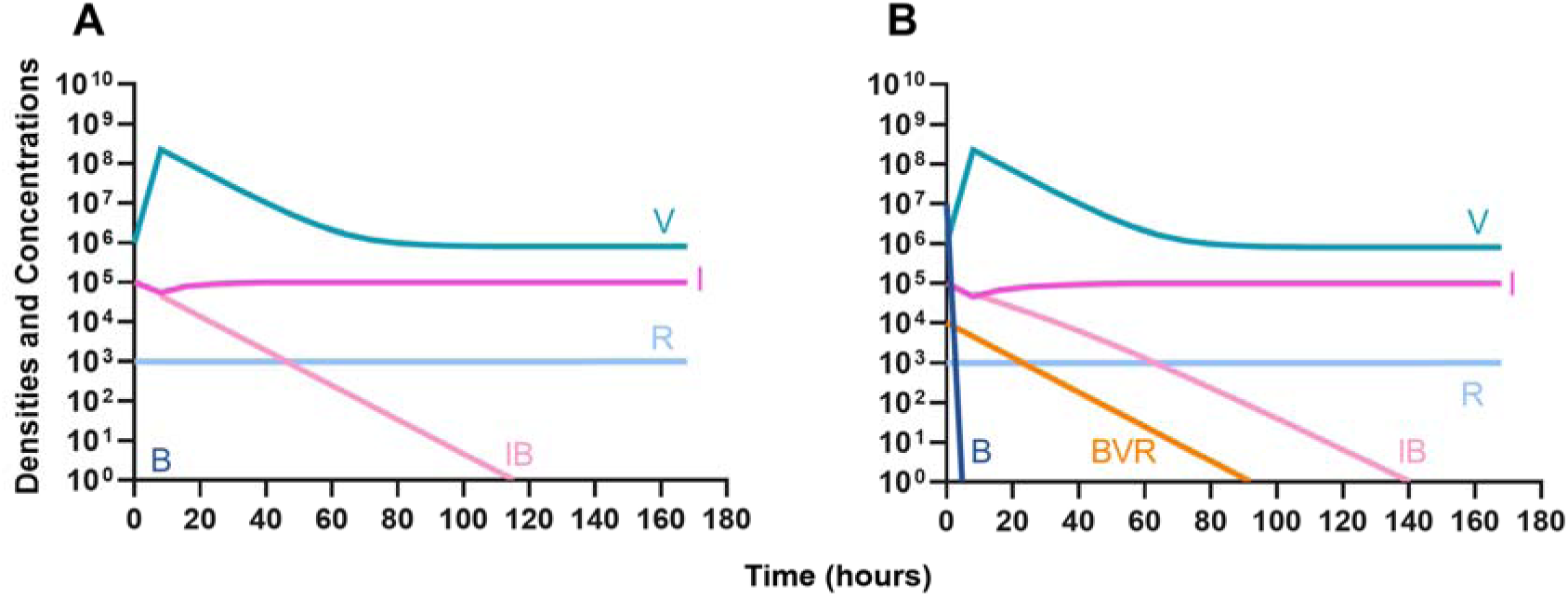
The interaction of bacteriophage and the innate immune system. **A.** A population of 1E7 bacteria when treated with phage in the presence of the immune system **C.** A population of 1E7 phage-sensitive bacteria and 1E4 phage-resistant bacteria treated with a phage in the presence of the immune system.

### Phage, antibiotics, and the innate immune system

We next expand our consideration of the joint action of treatment and the immune system to situations where phage and antibiotics are used in conjunction.

#### Bacteriostatic antibiotics

First, we evaluate the dynamics of infection when treated with a bacteriostatic drug and a lytic phage in the presence of the immune system. As above, we consider the effects that dosing order has on treatment outcome (Figure 11) and find that the effect of treatment dosing order is minimal, and all condition are capable of clearing the infection without the ascent of resistance. Although, the condition where the bacteriostatic drug is administered first does have the highest time to clearance (Figure 11B).

**Figure 11.**
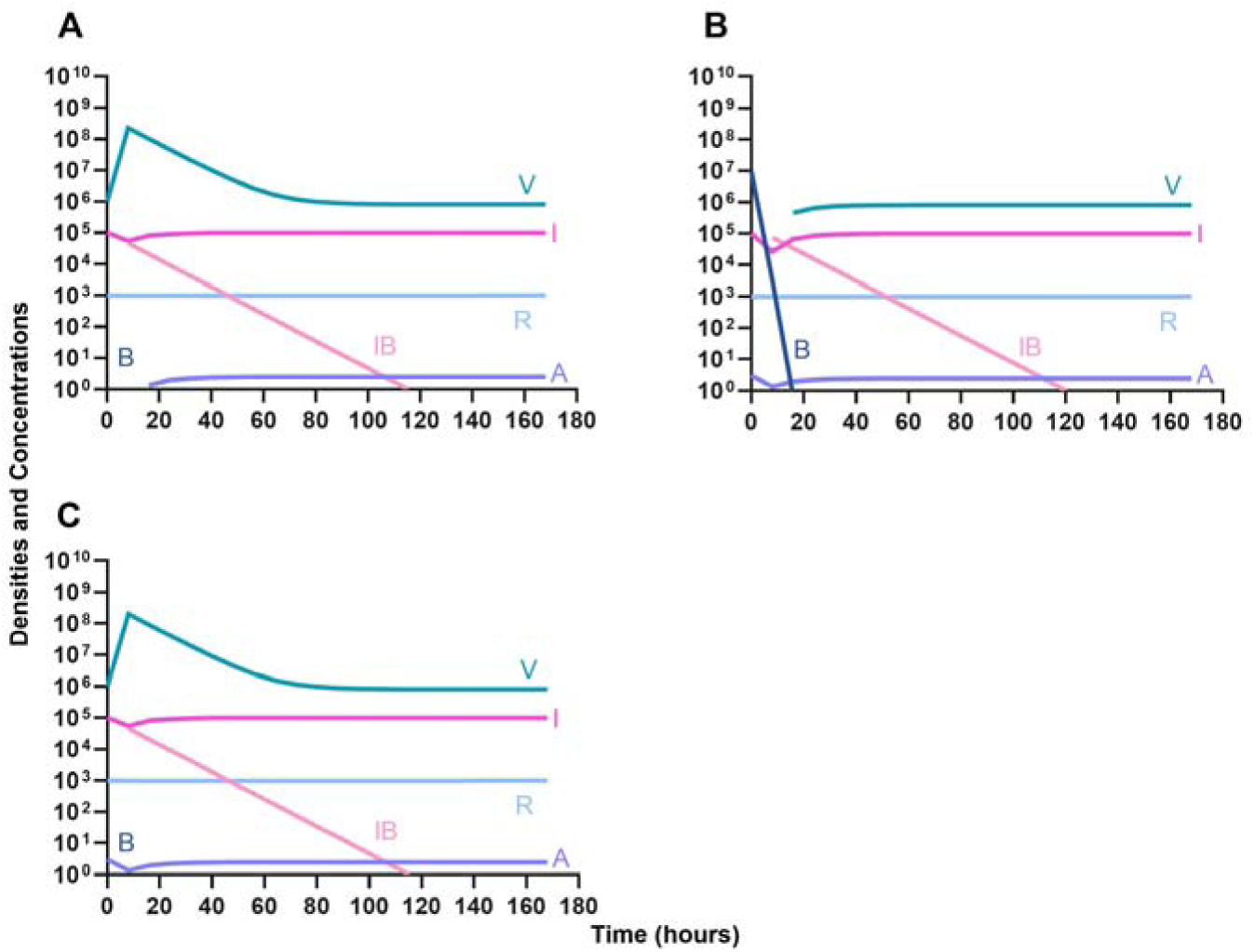
Treatment with a bacteriostatic antibiotic and a phage with differing dosing regimens in the presence of the immune system. **A.** A phage and then a bacteriostatic antibiotic **B.** A bacteriostatic antibiotic and then a phage **C.** Coadministration of a bacteriostatic antibiotic and a phage.

#### Bactericidal antibiotics

We determine the effect of dosing order for a bactericidal drug and phage with the innate immune system as in Figure 11. The results of these simulations in Figure 12 are parallel to those in Figure 11, once again demonstrating that there is no effect on treatment outcome with dosing order or using a bacteriostatic versus a bactericidal drug. However, the time to clearance is once again longer when the antibiotic is applied first (Figure 12B).

**Figure 12.**
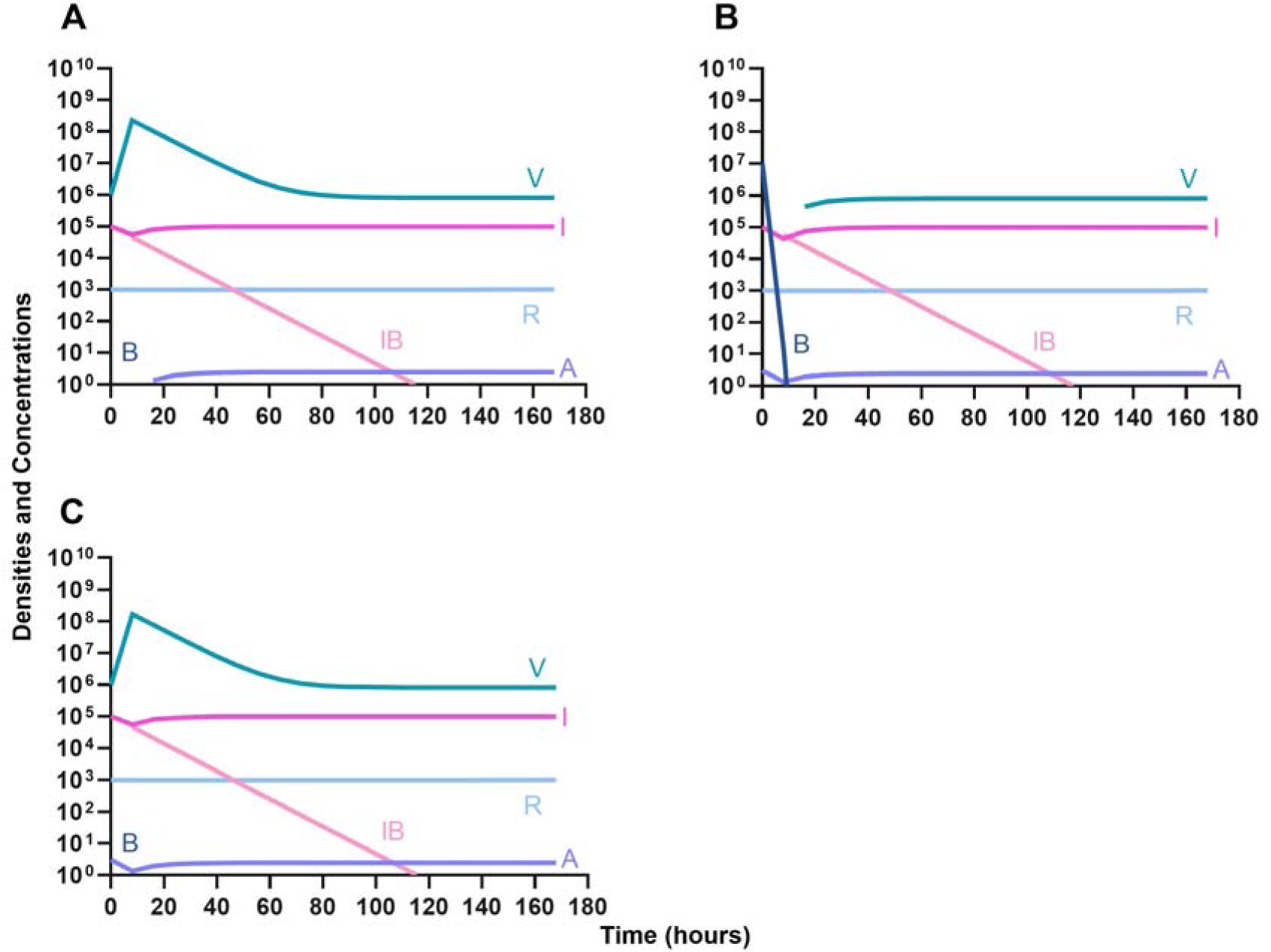
Treatment with a bactericidal antibiotic and a phage with differing dosing regimens in the presence of the immune system. **A.** A phage and then a bactericidal antibiotic **B.** A bacteriostatic antibiotic and then a phage **C.** Coadministration of a bactericidal antibiotic and a phage.

#### Suppression of resistance

Once again, the logic of using multiple treating agents is predicated upon the suppression of resistance. We finally consider three situations where a minor population is resistant to either one (Figure 13A and 13B) or both treating agents (Figure 13C). Ultimately, our results indicate that resistant subpopulations, regardless of what they are resistant to, will not ascend under treatment when the immune system is present. Interestingly, when a population that is resistant to both the phage and a bactericidal drug is initially dominant and at a very high density, treatment can still control and eventually clear the infection (Figure 13D).

**Figure 13.**
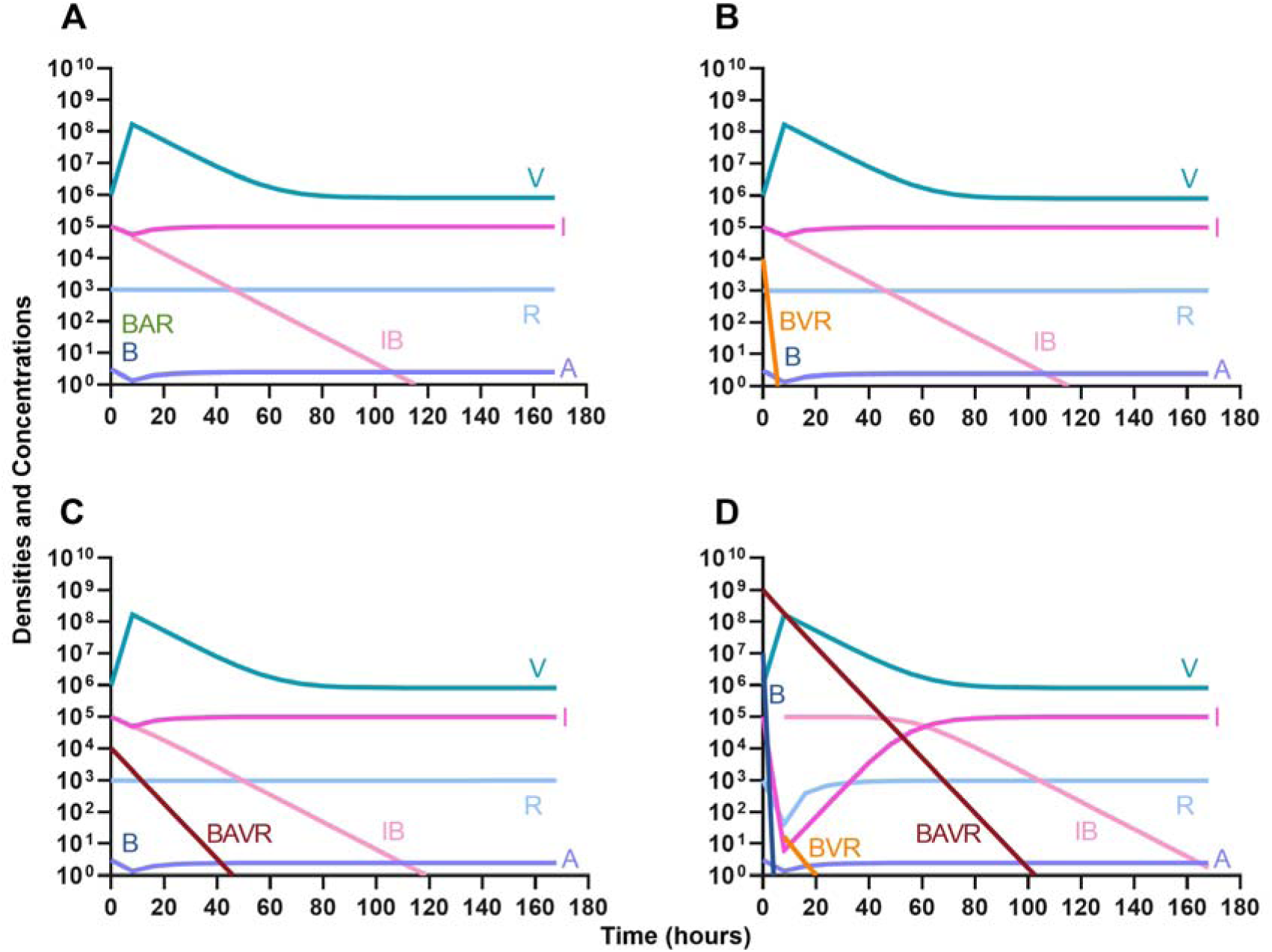
The ability of the immune system to suppress resistance. **A.** A population of 1E7 antibiotic-sensitive bacteria and 1E4 antibiotic-resistant bacteria treated with a bactericidal drug and a phage in the presence of the immune system **B.** A population of 1E7 phage-sensitive bacteria and 1E4 phage-resistant bacteria treated with a bactericidal drug and a phage in the presence of the immune system **C.** A population of 1E7 phage-sensitive and antibiotic-sensitive bacteria and 1E4 phage-resistant and antibiotic-resistant bacteria treated with a bactericidal drug and a phage in the presence of the immune system **D.** A population of 1E7 phage-sensitive and antibiotic-sensitive bacteria and 1E9 phage-resistant and antibiotic-resistant bacteria treated with a bactericidal drug and a phage in the presence of the immune system.

### A model of phage cocktails

Motivating the use of cocktails of phage, rather than a single phage, for treatment is the suppression (or elimination) of phage-resistant mutants. Here, we consider a model where treatment can be with up to three phages and bacteria resistant to each phage and the various combinations of the three phages can emerge. This model does not have the innate immune system, nor does it have antibiotics.

#### Single phage treatment

In the absence of the immune system, when a single phage is used for therapy, resistance to the treating phage very rapidly ascends to dominate and treatment fails (Figure 14). However, the phage is maintained over time due to the transition from the resistant state to the sensitive state.

**Figure 14.**
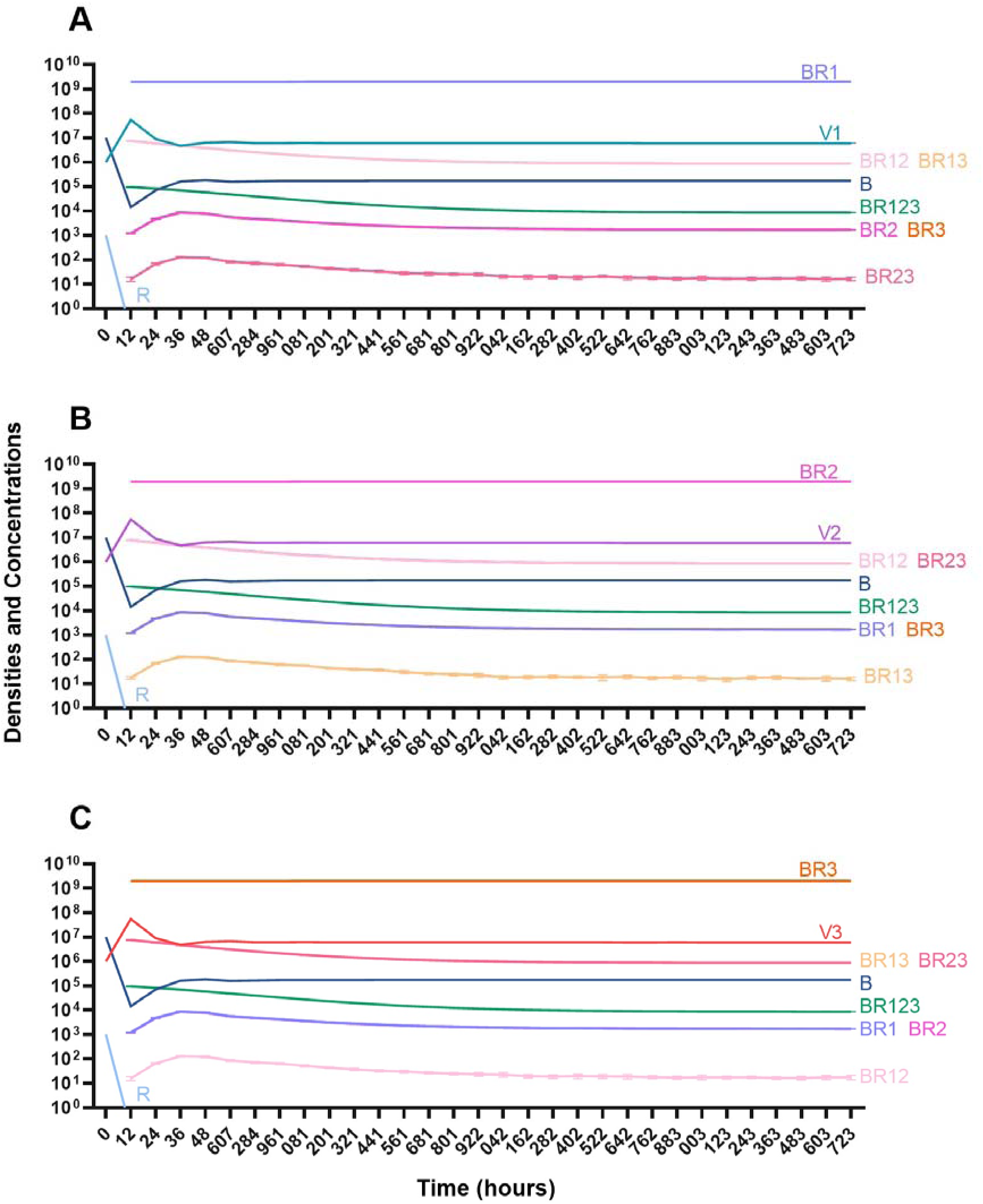
Treatment with a single phage. **A.** Treatment with Phage 1 **B.** Treatment with Phage 2 **C.** Treatment with Phage 3.

#### Two phage treatment

We then consider a situation where two phages are used in combination (Figure 15). Notably, the time before the mutant resistant to both treating phages ascends to dominance is longer than when one phage is used for treatment. However, since this model is stochastic, there is variability when the single-resistant mutants emerge and thereby variability in when the double-resistant mutants emerge.

**Figure 15.**
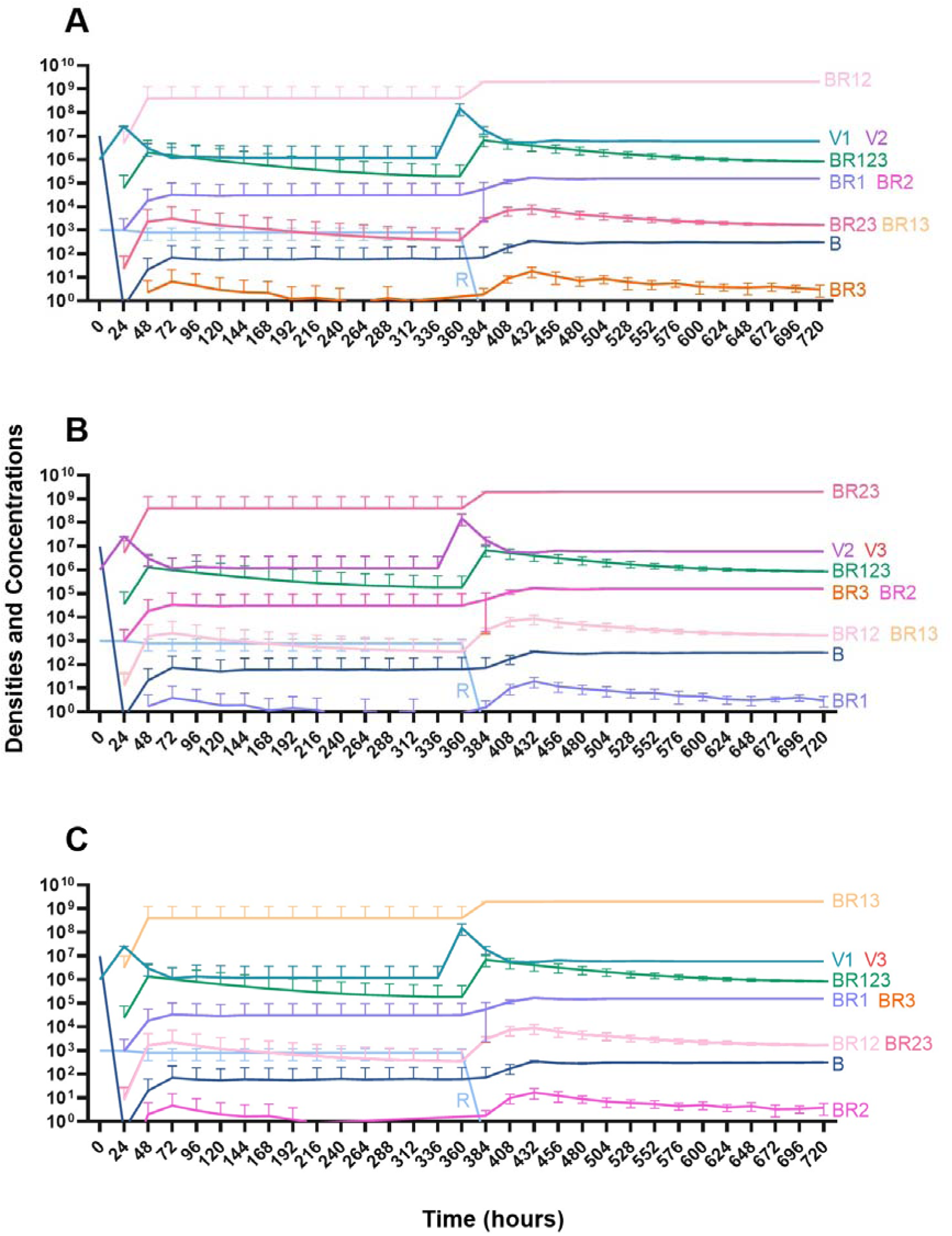
Treatment with two phages. **A.** Treatment with Phage 1 and Phage 2 **B.** Treatment with Phage 2 and Phage 3 **C.** Treatment with Phage 1 and Phage 3.

#### Three phage treatment

Finally, under treatment with three phages (Figure 16), single phage-resistant mutants arise at various times and give way to double-phage resistant mutants, before the triple-phage resistant mutants ultimately arise and dominate such that treatment fails, but it takes longer for treatment with three phages to fail compared to treatment with two phage and substantially long to fail that treatment with one phage.

**Figure 16.**
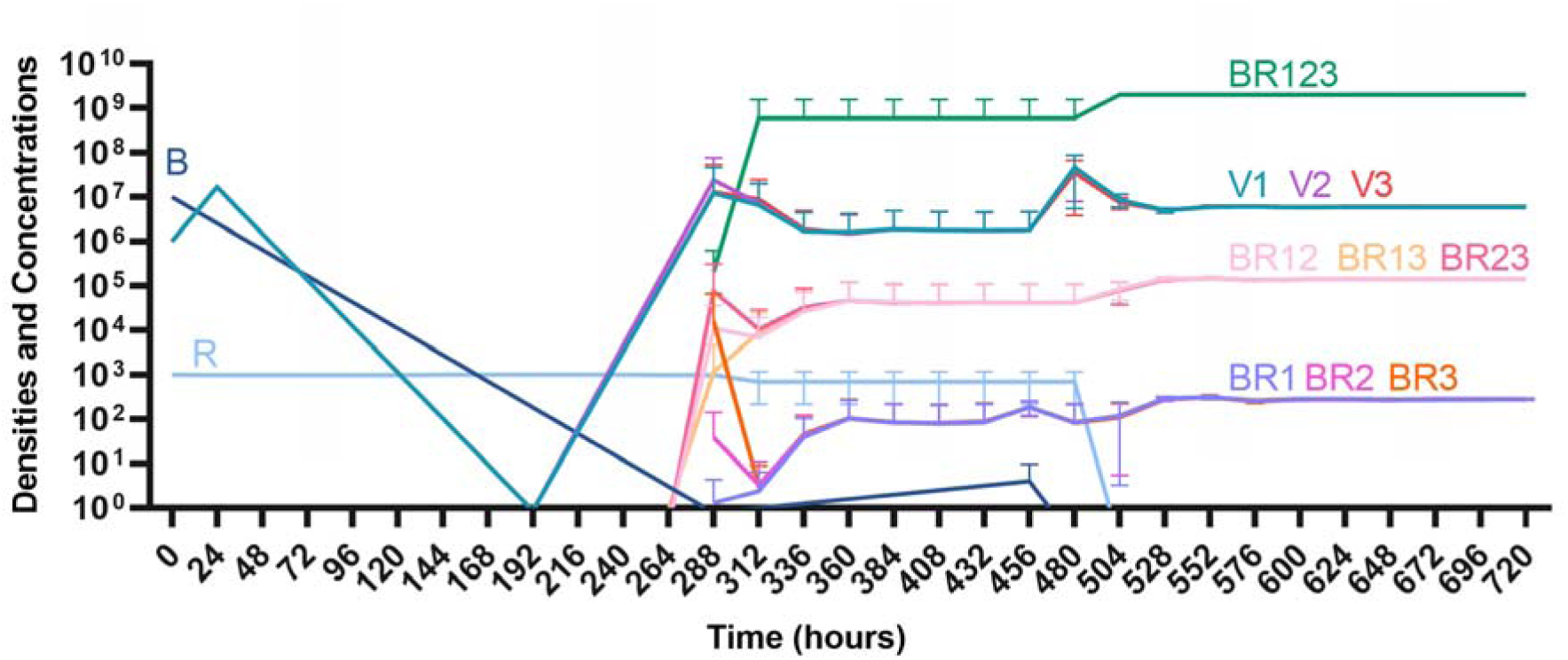
Treatment with three phages. Treatment with Phage 1, Phage 2, and Phage 3.

## Discussion

Motivated by the well warranted concern about the antibiotic-resistance crisis, there has been an increase in studies on the treatment of bacterial infections (24). There is no shortage of treatment options for infections given the numerous types and classes of antibiotics as well as burgeoning complementary therapies such as the use of bacteriophages (phages). However, many of the studies neglect the role of the host in the dynamics of infections, particularly the role of the innate immune system (25). To lay the foundation for further experimental studies, in this report, we create two mathematical and computer-simulation models that generate testable hypotheses about the population and evolutionary dynamics of bacterial infections under treatment with antibiotics and phage in the presence of the host’s innate immune system.

The results of the analysis of our models underscore the need to consider the role of the innate immune system in subsequent experimental studies. In the absence of the immune system, resistance to the treating agent invariable emerges independent of the treating agents or the regimens in which they are employed. On the other hand, when the immune system is present, resistance does not emerge; indeed, even when a high density of pan-resistant bacteria is present, the infection can still be controlled with treatment. As previously reported with numerous *in vitro* and *in vivo* studies, the difference in treatment outcome with bacteriostatic and bactericidal antibiotics is de minimis (17, 26).

The predictions of this theory are also congruent with previous results that demonstrate that phage can be as effective as antibiotics in controlling infections (27). This model also provides support for the intuitive conclusion that more phages are better than fewer phages. While resistance to multiple treating phages does ultimately emerge, the time for resistance to dominate for one treating phage is measured in hours, while the time for resistance to dominate for three treating phages is measured in days.

As with all purely theoretical studies, we have had to make assumptions about the parameters which we could not find in previous reports. One key parameter to which the model is incredibly sensitive which has not been estimated is the rate of phagocyte gobbling. For this report, we have elected to use a phagocyte gobbling rate constant that keeps the density of the infecting bacteria steady without the presence of any treatment. This assumption allows for us to determine the potential impact that the treatments and their order are having on the dynamics of the infection. However, these immune parameters, and moreover, all the parameters used in this study can be readily estimated experimentally.

Taken together, the analysis of our mathematical and computer-simulation models makes highly testable predictions about the dynamics of treatment which could be supported or rejected by using a mix of *in vitro* and *in vivo* models. It is the intent of these authors to explore the validity of the hypotheses generated above with the *Galleria mellonella* infection model system (28). Be that as it may, these predictions are agnostic to the experimental system and the hypotheses could easily be tested in other systems such as cell culture or mice (29, 30).

## Materials and Methods

### Numerical Solutions (Simulations)

For our numerical analysis of the coupled, ordered differential equations presented (Equations 1-12), we used Berkeley Madonna with the parameters presented in Table 2 (31). Copies of the Berkeley Madonna programs used for these simulations are available at www.eclf.net.

## Acknowledgements

We thank the other members of the Levin Lab for their comments on an earlier version of this manuscript.

## Funding Sources

BRL would like to thank the U.S. National Institute of General Medical Sciences for their funding support via R35GM136407 and the Emory University Antibiotic Resistance Center. The funding sources had no role in the design of this study and will not have any role during its execution, analysis, interpretation of the data, or drafting of this report. The content is solely the responsibility of the authors and does not necessarily represent the official views of the National Institutes of Health.

## Data Availability

The Berkeley Madonna programs used for these simulations are available at ECLF.net. All data are presented in this article or its supplementary materials.

## Notes

### Competing Interest Statement

The authors have declared no competing interest.

